## Supplemental Figure 1-5 for "Neural pathways and computations that achieve stable contrast processing tuned to natural scenes"

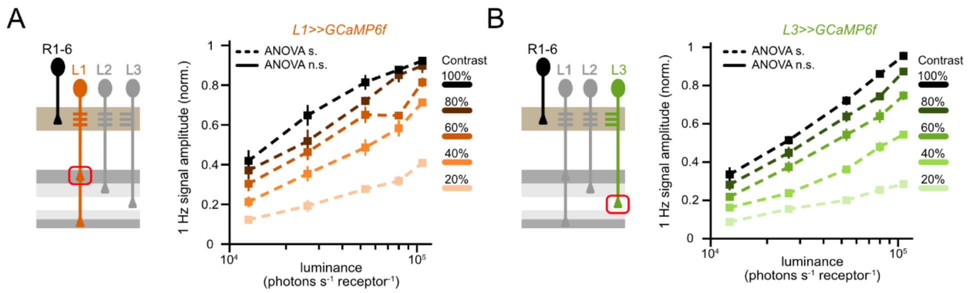


### Figure S1 related to Figure 2. L1 and L3 neurons do not exhibit luminance gain.

**(A-B)** Calcium imaging of L1 (orange), L3 (green) while stimulating the fly with drifting 1Hz gratings of changing contrast and luminance. **(A)** Mean contrast responses of L1 neurons across luminance, each curve represents a different grating contrast. One way ANOVA between the luminances for each contrast to assess luminance-invariant responses. Dashed lines represent $p<0.05$ and solid lines represent $p>0.05$. **(B)** Same as (A) for L3.

Error bars represent ±SEM. Means are calculated across flies. Sample sizes for (A-B), L1: n=5(36), L3: n=7(64). Sample sizes are given as: flies(cells).

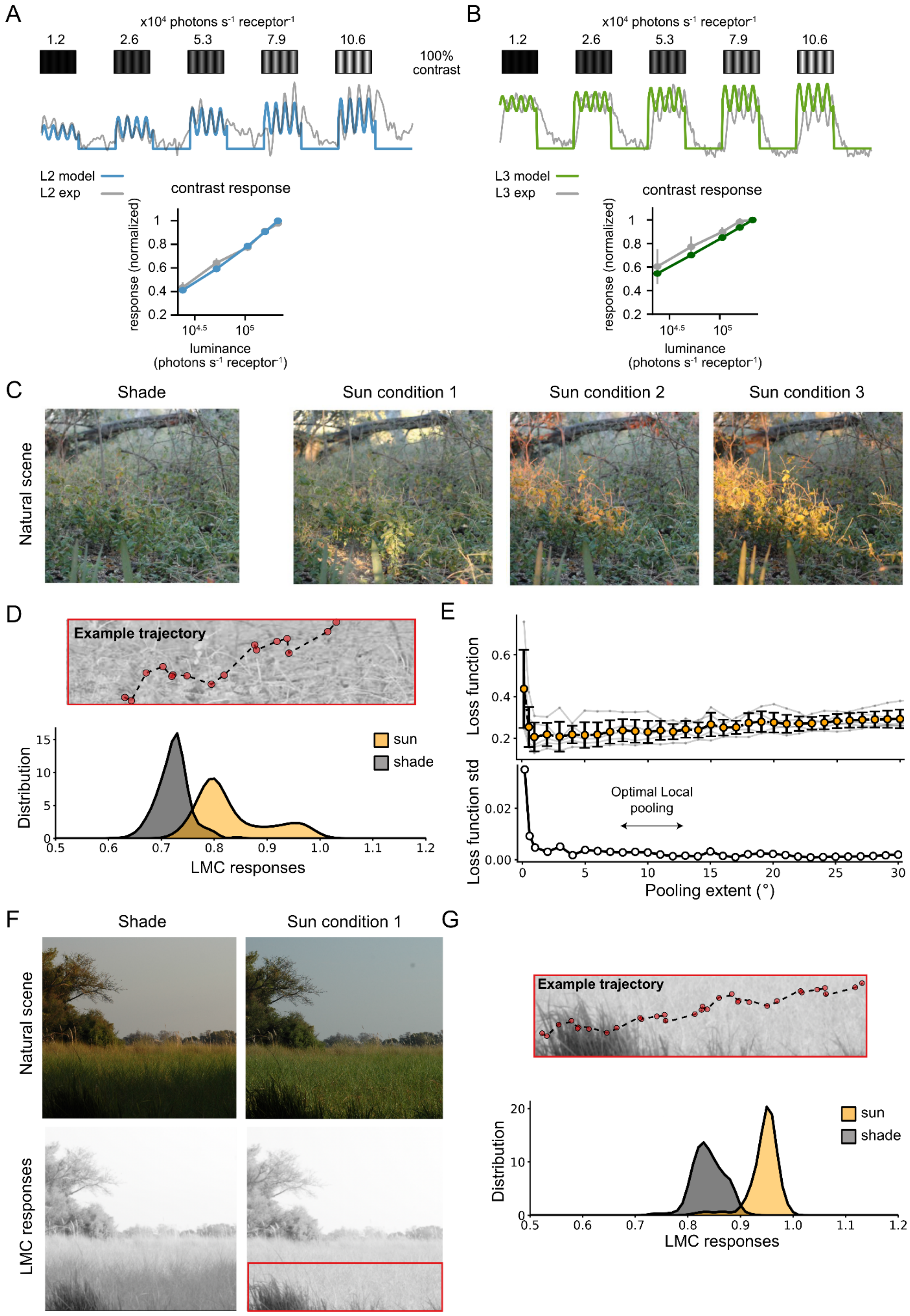


Figure S2 related to Figure 3. Simulating LMC responses with a model based on physiological data

**(A)** Left: Simulated L2 neuron responses (blue) based on a model fitted to two-photon *in vivo* calcium imaging data of L2 neurons (gray) while stimulating the fly with drifting 1Hz gratings of changing contrast and luminance (data from Figure 1). Right: Mean contrast responses (F1 amplitude) of the fitted L2 model (blue) and experimental data (gray) across different luminances.

**(B)** Same as (A) for L3 neurons.

**(C)** Natural scene as in Figure 3A under shade and three different sunny conditions. For the analysis of the cost function, we have used a total number of five sunny conditions.

**(D)** Above: Example of a single navigation trajectory as in Figure 3C with a different orientation. We used a total number of 15 trajectories to sample LMC responses. Below: Probability distribution of LMC contrast responses in both shaded and sunny conditions for a total number of 15 simulated stochastic trajectories with the same orientation and direction. The distributions of LMC responses under both luminance conditions change depending on the sample trajectory conditions, e.g. orientation. The overlap between both distributions continues to be small.

**(E)** Above: Loss function (Methods) as a function of pooling extent. We calculate the loss function using five different sunny conditions captured from the same natural scene; here represented by the gray traces. Orange markers and error bars correspond to the mean and standard deviation over these five luminance conditions. Below: Standard deviation of the loss function as a function of pooling extent. The region in yellow highlights the pooling extent region that optimizes contrast encoding by minimizing both the mean and standard deviation (std) of the loss function.

**(F)** Second natural scene, as in Figure 3G, for both shade and sunny conditions with the corresponding LMC responses. The red square shows the visual location where the navigation trajectory to sample LMC responses was simulated.

**(G)** Above: Example of a single navigation trajectory for the second natural scene shown in F and in Figure 3G used to sample contrast responses of LMC neurons under dynamic conditions. Below: Probability distribution of LMC contrast responses in both shaded and sunny conditions for a total number of 15 simulated stochastic trajectories with the same orientation and direction.

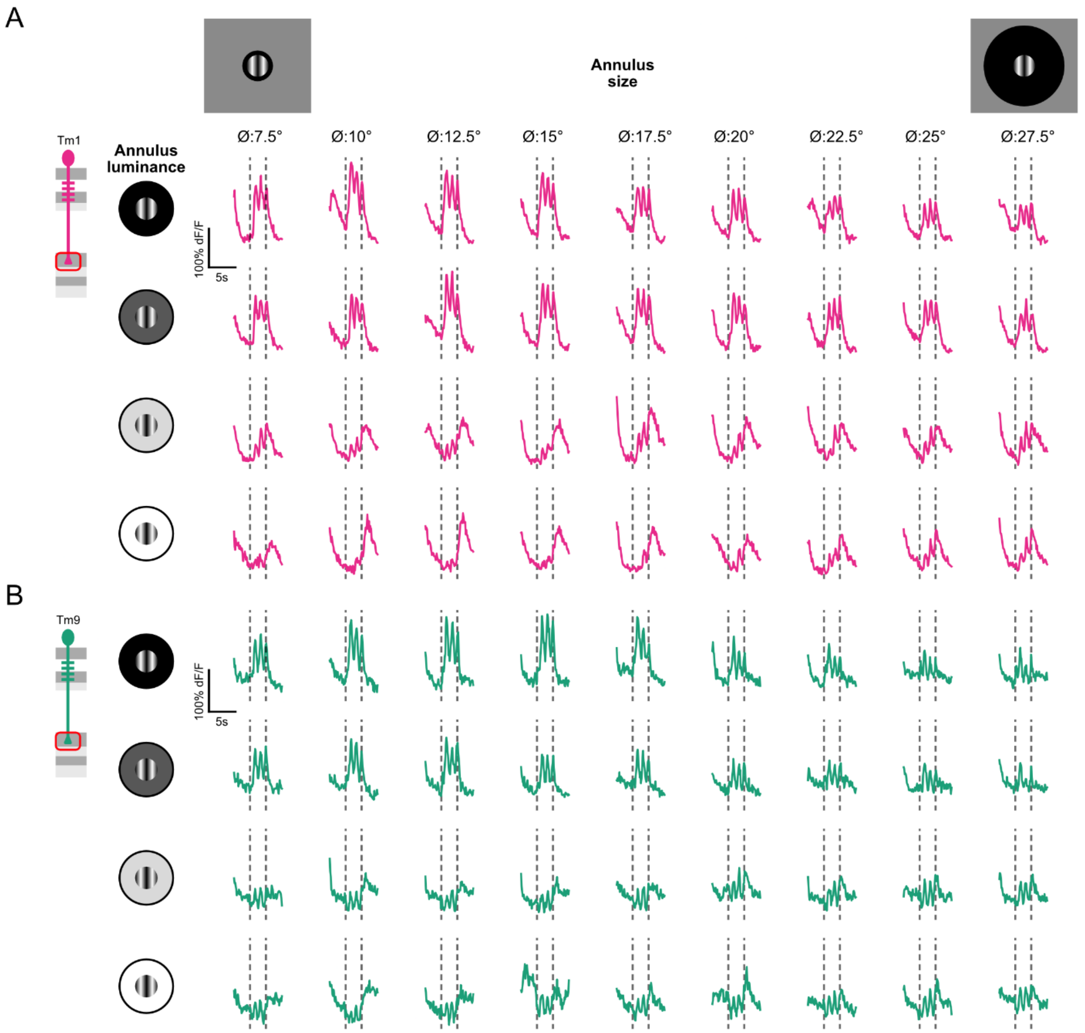


Figure S3 related to Figure 4. Tm1 and Tm9 example responses to centered gratings with changing background annulus

**(A)** Example calcium responses of a single Tm1 axon terminal (magenta) to the stimulus with a circular 5° sinusoidal grating moving at 1Hz with constant contrast (100% Michelson) and constant luminance. The circular grating contained background annulus of changing outer circle diameters (7.5° to 27.5°) and changing luminances.

.

**(B)** Same as (A) for Tm9 axon terminals (teal).

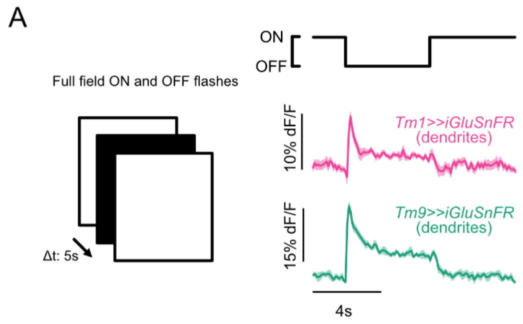


#### Figure S4 related to Figure5. Tm1 and Tm9 neurons receive wide glutamatergic inputs

**(A)** Mean glutamate responses of Tm1 (magenta) and Tm9 (teal) neurons to alternating ON ($I_{max}$) and OFF full field flashes. $I_{max}$= $2.17x10^5$.

Error bars and patches represent ±SEM. Means are calculated across flies. Sample sizes for (A), Tm1>>iGluSnFR n=17(224), Tm9>>iGluSnFR n=7(82). Sample sizes are given as: flies(cells).

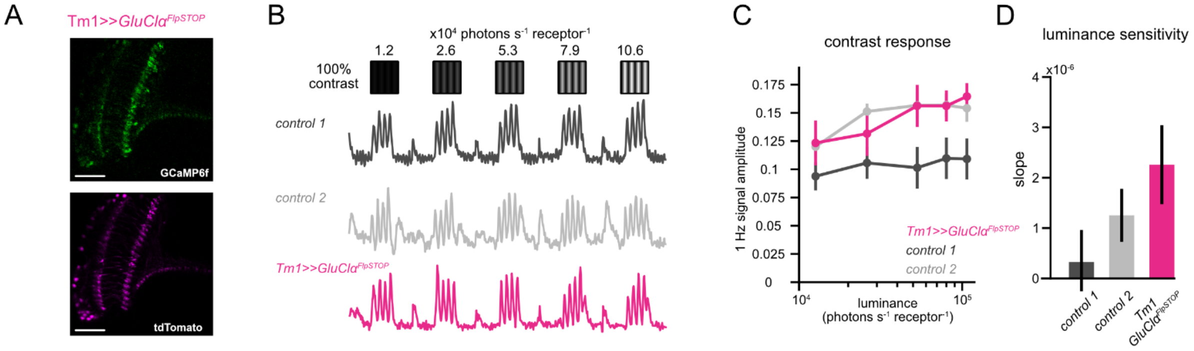


#### Figure S5 related to Figure 6. GluClα is not involved in luminance gain in Tm1

**(A)** tdTomato expression (magenta) confirms *GluClα*$^{FlpSTOP}$ in Tm1 neurons. Scale bars are 50μm.

**(B)** Calcium imaging of *Tm1>>GluClα*$^{FlpSTOP}$ (magenta) and control genotypes (black: no Flp, and gray: heterozygous control) while stimulating the fly with drifting 1Hz gratings of constant 100% Michelson contrast and changing luminances and representative calcium responses of a single axon terminal for each genotype.

**(C)** Mean contrast responses (F1 amplitude) of each genotype across luminance.

**(D)** Slopes of the contrast responses depicting the luminance dependence for each neuron. $^{**}p<0.01$, $^{***}p<0.001$, one way ANOVA with post-hoc Tukey HSD test.

Error bars represent ±SEM. Means are calculated across flies. Sample sizes for (C), control 1: 6(75), control 2: n=6(57), *Tm1>>GluClα*$^{FlpSTOP}$: n=10(131). Sample sizes are given as: flies(cells).
